# Validation of Salivary Oxytocin and Vasopressin as Biomarkers in Domestic Dogs

**DOI:** 10.1101/151522

**Authors:** Evan L. MacLean, Laurence R. Gesquiere, Nancy Gee, Kerinne Levy, W. Lance Martin, C. Sue Carter

## Abstract

**Background:** Oxytocin (OT) and Vasopressin (AVP) are phylogenetically conserved neuropeptides with effects on social behavior, cognition and stress responses. Although OT and AVP are most commonly measured in blood, urine and cerebrospinal fluid (CSF), these approaches present an array of challenges including concerns related to the invasiveness of sample collection, the potential for matrix interference in immunoassays, and whether samples can be collected at precise time points to assess event-linked endocrine responses.

**New Method:** We validated enzyme-linked immunosorbent assays (ELISAs) for the measurement of salivary OT and AVP in domestic dogs.

**Results:** Both OT and AVP were present in dog saliva and detectable by ELISA and high performance liquid chromatography – mass spectrometry (HPLC-MS). OT concentrations in dog saliva were much higher than those typically detected in humans. OT concentrations in the same samples analyzed with and without sample extraction were highly correlated, but this was not true for AVP. ELISA validation studies revealed good accuracy and parallelism, both with and without solid phase extraction. Collection of salivary samples with different synthetic swabs, or following salivary stimulation or the consumption of food led to variance in results. However, samples collected from the same dogs using different techniques tended to be positively correlated. We detected concurrent elevations in salivary and plasma OT during nursing.

**Comparison with Existing Methods:** There are currently no other validated methods for measuring OT/AVP in dog saliva.

**Conclusions:** OT and AVP are present in dog saliva, and ELISAs for their detection are methodologically valid.

## Introduction

Oxytocin (OT) and arginine vasopressin (AVP) are phylogenetically conserved neuropeptides with wide-ranging effects on social behavior, cognition, anxiety, and stress responses (Carter, 1998; Carter, Grippo, Pournajafi-Nazarloo, Ruscio, & Porges, 2008; Donaldson & Young, 2008). Although the behavioral effects of OT and AVP are diverse, and depend on the species (Insel & Shapiro, 1992; Kramer, Cushing, Carter, Wu, & Ottinger, 2004), site of action in the brain (Kelly & Goodson, 2014), and individual and contextual factors (Bartz, Zaki, Bolger, & Ochsner, 2011), both peptides have been highly implicated in the processes of bond formation and social attachment in numerous species (Carter, 1998). Based on these associations, scientists have begun to explore whether these peptides may play similarly important roles in social interactions between species, for example between humans and domestic dogs (Beetz, Uvnäs-Moberg, Julius, & Kotrschal, 2012; Carter & Porges, In press; MacLean & Hare, 2015; Thielke & Udell, 2015).

Although only a handful of studies have investigated the roles of OT in human-animal interaction (HAI), early findings support the hypothesis that OT both facilitates, and responds to key processes in HAI. For example, both dogs and humans exhibit an increase in OT (measured in blood or urine) following affiliative social interaction (Nagasawa, Kikusui, Onaka, & Ohta, 2009; Nagasawa et al., 2015; Odendaal & Meintjes, 2003; Rehn, Handlin, Uvnäs-Moberg, & Keeling, 2014). Increases in urinary OT have also been detected following other (presumably) pleasurable experiences in dogs, including eating and exercise, suggesting that OT may generally index positive emotions, providing a potentially useful biomarker for assessing welfare (Mitsui et al., 2011). Additionally, dogs treated with intranasal OT have been documented to exhibit increased affiliative behavior toward humans and other dogs (Romero, Nagasawa, Mogi, Hasegawa, & Kikusui, 2014, 2015), as well as increased ‘optimism’ about ambivalent stimuli (Kis, Hernádi, Kanizsár, Gácsi, & Topál, 2015). Several recent studies also reveal that administration of intranasal OT may enhance dogs’ sensitivity to cooperative-communicative signals (Macchitella et al., 2016; Oliva, Rault, Appleton, & Lill, 2015), an aspect of dog cognition that may be evolutionarily convergent with humans (Hare, 2016; Hare & Tomasello, 2005; MacLean, Herrmann, Suchindran, & Hare, In Press), and which likely plays important roles in dog-human relationships.

Currently we know less about the role of AVP in dog behavior, however, findings from other mammalian species suggest that this neuropeptide makes similarly important contributions to social behavior (Caldwell, Lee, Macbeth, & Young III, 2008). To our knowledge there have only been two studies exploring the relationship between AVP and behavior in dogs. Hydbring-Sandberg et al. (2004) report that following exposure to potentially fear-inducing stimuli, plasma AVP concentrations were positively associated with the extent of fearful reactions. Second, in a case-control study, MacLean et al. (Submitted) show that dogs with a history of aggression toward conspecifics where characterized by lower free, but higher total AVP, than age-, sex- and breed-matched controls. Therefore, preliminary findings are consistent the notion that AVP is primarily involved in upregulation of the hypothalamic-pituitary-adrenal (HPA) axis, and the genesis of anxious, fearful, or aggressive behavioral states (Neumann & Landgraf, 2012).

Despite progress investigating the role of these neuropeptides in dog behavior and cognition, current methods for assessing endogenous OT and AVP are restricted to measures derived from urine or blood, which present a host of challenges related to design, analysis, and welfare. For example, urinary samples capture wide windows of hormonal activity, and the process of urine collection can be logistically challenging. While blood samples may provide greater temporal resolution and sensitivity to acute changes in peptide release, blood draws are invasive, potentially stressful, and peptide binding in blood can interfere with measurement (Brandtzaeg et al., 2016; Martin & Carter, 2013; Martin, 2014). In contrast, salivary measures are comparatively noninvasive (relative to blood draws) and can be collected at precise time points to assess event-linked endocrine responses (which is considerably more challenging with urine samples).

Several human studies have employed salivary measures of OT and AVP, yet there is limited consensus about the validity of this approach. Horvat-Gordan et al. (2005) conducted a series of methodological studies and concluded that OT was not a valid biomarker in human saliva. These researchers determined that the levels of OT in human saliva were very low, and salivary samples yielded poor parallelism when diluted against the standard curve. In addition to these empirical results, Horvat-Gordan et al. (2005) provided additional theoretical arguments why OT was unlikely to be a valid biomarker in saliva. For example, the molecular weight of OT may restrict transport to saliva, and given that OT is commonly detected at low levels in blood, the concentrations in saliva may be even lower, and below the limit of detection for most current platforms. However, with regard to the latter concern, recent findings suggest that OT concentrations in blood are much higher than many have assumed, but may evade detection due to binding to plasma proteins (Brandtzaeg et al., 2016; Martin & Carter, 2013). Other researchers have reported greater success measuring OT in saliva, including good technical characteristics in methodological studies (Carter et al., 2007), and sensitivity to OT-regulated biological processes, such as lactation (White-Traut et al., 2009).

To our knowledge there have been no studies evaluating the feasibility of measuring of OT or AVP in dog saliva. However, given that dog salivary samples are commonly used for the measurement of other hormones, such as cortisol or testosterone (Cobb, Iskandarani, Chinchilli, & Dreschel, 2016; Dreschel & Granger, 2016), validated methods for measuring OT and AVP in dog saliva could provide a versatile and noninvasive approach for future research on the roles of OT and AVP in dog behavior and cognition, as well as dog-human interaction. Here, we report a series of studies investigating the potential of OT and AVP as salivary biomarkers in dogs.

## Methods

### Subjects

Biological samples were collected from client-owned pet dogs at the Duke Canine Cognition Center (DCCC; Durham, NC, USA) and from dogs in the assistance dog breeding colony at Canine Companions for Independence (Santa Rosa, CA, USA). The pet dog sample consisted of 5 dogs (female Labrador retriever, 4 years; male Yorkshire terrier, 9 years; female border collie, 2 years; male border collie, 9 years, female springer spaniel, 9 years). All pet dogs lived and were cared for in human homes during the study, and saliva samples were collected during short visits to the DCCC (~10 minutes), following client consent. All pet dogs were familiar with this environment through previous visits for participation in behavioral research studies. For the comparison of sample collection techniques, we collected repeated samples (see method) from 20 assistance dogs in training at CCI (9 male, 11 female; 1 golden retriever, 4 Labrador retrievers, 15 Labrador X golden crosses, mean age = 1.83 years, SD = 0.24 years). These dogs were pair-housed in indoor-outdoor kennels and saliva samples were collected in the dog’s kennel. Lastly, for our nursing studies we collected samples from 6 dams (all Labrador-golden crosses, mean age = 2.98 years, SD = 1.11 years) at CCI’s Canine Early Development Center. Client consent was obtained for participation of all CCI dogs and all animal procedures were approved by the Duke University IACUC (protocol #’s: A303-11-12 & A138-11-06).

### Sample Collection Techniques

Dog saliva samples are routinely collected using absorbent swabs which can be placed inside the dog’s mouth to passively absorb saliva (Dreschel & Granger, 2009). To evaluate the effects of collecting samples with different swabs, artificially stimulating salivary production, or eating prior to sample collection, we collected multiple samples from the same subjects (N = 20) using a variety of procedures. We evaluated two commercially available swabs (SalivaBio Children’s Swab, Sarstedt Salivette® – Cortisol). Although both swabs are constructed of a synthetic material, the Salimetrics swab is thinner, more pliable, and designed to passively absorb saliva without chewing, whereas the Sarstedt swab has a larger diameter, is more rigid, and is designed to be chewed on to absorb saliva. To limit the risk of swabs being swallowed, we prevented dogs from chewing swabs and instead placed the swab between the mandibular teeth and cheek for ~1 minute, while gently holding the dog’s mouth closed. Following sample collections dogs were rewarded with a small treat. Although we were able to obtain saliva using both swabs, the Children’s Swab absorbed larger amounts of saliva, and fitted more comfortably in the dog’s cheek. Therefore, coupled with the fact that this swab has been validated for a larger range of analytes, we opted to use the Children’s Swab for all subsequent sample collections. However, we report a comparison of results from both swabs below.

To explore the effect of salivary stimulation/saliva flow rate, we used 125 μl of a solution of citric acid dissolved in water (0.17 g/mL) which was spritzed into the dog’s mouth using a spray bottle (the solution had a pH of 3.0, acidity comparable to that of fresh lime or grapefruit). Saliva samples were collected using the Children’s Swab 30 seconds following the application of citric acid. Lastly, to explore whether eating prior to sample collection interfered with results, dogs were fed four pieces of Eukanuba™ Large Breed kibble, and we collected samples using the Children’s Swab 30 seconds after dogs consumed the kibble.

### Sample processing procedures

To determine if dog saliva samples should be extracted prior to analysis we generated pools of saliva samples collected from dogs at the Duke Canine Cognition Center, processed these pools according to a number of different protocols (described below), and used the resulting samples to assess parallelism, accuracy/recovery, and intra- and inter-assay coefficients of variation for the ELISAs. Samples for these pools were collected with Children’s Swabs, cut into two pieces, and placed between the mandibular teeth and cheek for ~1 minute (one swab on each side of the mouth). Following sample collections dogs were rewarded with a small treat and samples were immediately frozen at -20°C. Following this initial freeze cycle, samples were thawed and centrifuged at 3,000 *g* for 20 minutes at 4°C. Individual samples were then pooled, vortexed thoroughly, and divided into 1 mL aliquots that were frozen at − 20°C.

For solid phase extraction (SPE) samples were diluted 1:2 with 0.1% trifluoroacetic acid (TFA), and centrifuged at 15,000 RPMs for 15 minutes. OT and AVP saliva samples were run on separate Oasis PRiME cartridges (Waters Corporation) which were conditioned with 1 mL acetonitrile (ACN), followed by 1 mL of 0.1% TFA in H_2_O before loading the sample (gravity-fed). Cartridges were then washed with 6 mL of 0.1% TFA in H_2_O. Lastly, AVP samples were eluted using 90% ACN, 0.1% TFA, and OT samples were eluted using 95% ACN, 0.1 % TFA. All samples were then evaporated to dryness under a steady stream of air at 37°C, and frozen until assay.

### ELISA kits

For measurement of OT we used commercially available kits from Arbor Assays (K048) and Cayman Chemical (Item #500440). We initially compared accuracy, linearity and parallelism with both kits, but all subsequent analyses were performed with the Cayman Chemical kit based on better spike recovery and parallelism using this kit (see below). AVP was measured using a commercially available kit from Enzo Life Sciences (ADI-900-017A).

### Parallelism, Accuracy and Coefficients of Variation (CVs)

To evaluate linearity and parallelism, a pool of dog saliva was measured at a range of dilutions between 100-10% of its fully concentrated value. Linearity of sample dilutions was assessed by running linear models with the observed value predicted by the expected value at each dilution. Parallelism was assessed by plotting the log_10_ expected concentrations of samples as a function of logit of the proportion binding [B/B0] (these transformations allow the normally sigmoidal shape of the standard curve to be presented linearly). Following the method developed by Plikaytis et al. (1994) we calculated CVs for the corrected sample concentrations across dilutions as a statistical measure of parallelism. Parallelism was considered to be acceptable if these CVs were less than or equal to 20% (Plikaytis et al., 1994). Accuracy was measured by mixing 50 μl of pooled samples with 50 μl of kit standards and calculating the observed vs. expected values for these samples (expected = ½ standard value + ½ sample value). Intra-assay CVs were calculated by running 10 replicates of the same sample during the same assay and inter-assay CVs were calculated by running the same control samples across multiple assays (OT: N = 8, AVP: N = 7).

### Comparison of ELISA and HPLC-MS

Twenty samples were analyzed with and without extraction using ELISA as well as high performance liquid chromatography – mass spectrometry (Martin Protean).

### Biological validation: salivary OT during nursing

Because OT has long been known for its role in lactation, as a biological validation we sought to determine whether measures of salivary OT would be sensitive to the acute release of oxytocin associated with milk let-down. We collected blood and saliva samples from 6 dams at baseline and during nursing (blood samples were not possible with one dam and only saliva was collected in this instance). For two dams we also obtained post-nursing blood and saliva samples. Blood samples (3 mL) were collected via cephalic venipuncture using a winged blood collection set and 3 mL EDTA Vacutainer®. For the two dams from which we collected three samples, the third sample was collected from the jugular vein. We opted not to use a catheter for repeated samples because previous experience with that approach appeared to have greater potential for inducing stress in the dog than individual blood draws (e.g., due to the prolonged period in which the catheter is inserted, continuous wrapping/unwrapping/flushing of the line, and repositioning if venous access was lost).

Prior to sample collection, dams were separated from their litters for at least 30 minutes to ensure that no nursing took place immediately prior to the baseline samples. Following this pre-test period, we collected a baseline blood and saliva sample from the dam, and then reunited her with the litter. Once the majority of puppies began active suckling we collected a second blood and saliva sample (as nursing continued). When nursing concluded (~10 minutes later) we collected a final blood and saliva sample. All nursing plasma OT samples were processed using solid phase extraction procedures previously validated in our lab (MacLean et al., Submitted), and saliva samples were analyzed without extraction based on the results of our methodological validation.

### Statistical Analysis

Statistical analyses were performed in the R environment for statistical computing (Team, 2016). To assess associations between results from different detection methods, or between different sample preparations (with or without extraction) we used Pearson correlation with log transformed data. In cases where results between different methods were highly correlated, we used Bland-Altman analyses to inspect (Bland & Altman, 1986) biases between methods. To compare concentration values from the same samples analyzed with and without extraction we used paired-sample t-tests. To compare samples collected from the same dog using different techniques (e.g. swab type, salivary stimulation) we used linear mixed models with a fixed effect for sample collection procedure and a random effect for subject ID. Following significant main effects we used Tukey’s HSD tests for post-hoc comparisons.

For parallelism analyses, some data points near the lower limits of detection (a less reliable region of the standard curve) deviated from an otherwise linear relationship between the remainder of the observations. To inspect these cases, we calculated hat values for all data points in the linear model. Observations with hat values greater than 2x the mean hat value were treated as outliers, and the model was refit without these points (N = 2, in both cases the lowest concentration data point in the series of dilutions). To assess changes in salivary and plasma OT during nursing we used linear mixed models with a fixed effect for time point (baseline, nursing) and a random effect for dam ID. All linear models were initially fitted using raw (untransformed) data. In all cases, we inspected model residuals, and refit the model using log transformed data if the dispersion of residuals increased as a function of the predicted values (this occurred for only one model: the comparison samples collected using different techniques). All inferential tests used a symmetrically-distributed alpha value of 0.05.

## Results

### OT and AVP concentrations in dog saliva

Using ELISA, both OT and AVP were detectable in dog saliva regardless of whether or not samples were extracted. However, non-extracted samples yielded considerably higher concentrations. OT levels in non-extracted dog saliva (N = 20) had a median value 258 pg/mL (range = 207-471 pg/mL) with the Arbor Assays kit, and 679 pg/mL (range = 356-1073 pg/mL) with the Cayman Chemical kit. Extracted OT salivary samples (N = 20) had a median value of 41 pg/mL (range = 27-105 pg/mL) with the Arbor Assays kit, and 260 pg/mL (range = 181-418 pg/mL) with the Cayman Chemical kit. Non-extracted samples yielded significantly higher values than extracted samples with both kits (Cayman: t_19_ = -14.87, p < 0.01, Arbor: t_18_ = -20.76, p < 0.01). However, within both kits analyses of the same samples with and without extraction were highly correlated (Arbor: r = 0.80 p < 0.01; Cayman r = 0.61, p < 0.01). Bland-Altman analyses revealed that non-extracted samples yielded concentrations that were, on average, 428.7 pg/mL (Cayman Chemical) and 228.6 pg/mL (Arbor Assays) higher than extracted samples. However, the bias between these methods varied as a function of the mean OT concentration. Linear models predicting the difference between the two measures as a function of their mean revealed significant positive slopes with both kits (Cayman: β = 1.08, t_18_ = 7.25, p < 0.01; Arbor: β = 0.94, t_17_ = 7.56, p < 0.01). Despite within-kit correlations, samples measured using the same sample preparation protocol (e.g. extracted or not) were not strongly correlated between the Arbor Assays and Cayman Chemical kits (extracted:r = 0.18, p = 0.45; non-extracted: r = 0.08, p = 0.73).

AVP concentrations in non-extracted saliva had a median value of 454 pg/mL (range = 228-1489 pg/mL), but extracted samples yielded significantly lower concentrations (median = 5 pg/mL, range = 2-11 pg/mL, t_18_= −7.01, p < 0.01), and required concentration during the extraction procedure to be detectable within a reliable region of the kit’s range. However, even with twofold concentration, many samples fell near the limit of detection, and outside an ideal region of the standard curve. Unlike salivary measures of OT, there was no correlation between the same samples analyzed for AVP with and without extraction (r = 0.02, p = 0.94). However, as noted above, many extracted salivary AVP samples were near the lower limit of detection and assay precision may be poor in this range.

Analysis by HPLC-MS confirmed the presence of both OT and AVP in dog saliva. For OT, 9 of 20 samples were below the limit of detection (~2 pg/mL) and the remaining samples had a median concentration of 18 pg/mL (range = 8-49 pg/mL). For AVP, 5 of 20 samples were below the limit of detection (~ 2 pg/mL) and the remaining samples had a median concentration of 25 pg/mL (range = 5-73 pg/mL). For comparison between HPLC-MS and ELISA we set all samples that were below the HPLC-MS limit of detection to the lower limit of detection (2 pg/mL). For OT, all ELISA methods (extracted and non-extracted samples analyzed with both kits) exhibited a positive correlation with HPLC-MS (mean r = 0.31, however only one of these correlations was significant (Arbor extracted, r = 0.45, p = 0.05). For AVP, neither ELISA measure yielded concentrations that were correlated with those obtained from HPLC-MS (average r = -0.23).

### Parallelism, Accuracy and Coefficients of Variation (CVs)

Figure 1 shows regressions of observed peptide concentrations predicted by the expected value across a series of dilutions with extracted and non-extracted saliva. In all cases dilutions were linear, had slopes close to the expected value of 1, and the dilution factor explained the vast majority of variance in detected concentrations. Parallelism data are shown in Figure 2. For OT samples analyzed using the Arbor Assays kit there were mild deviations between the slopes from the samples and standards regardless of the sample preparation protocol. Across all dilutions the CVs were 25.3% for non-extracted samples, and 31.9% for extracted samples. However, in both cases parallelism was best in the highest range of concentrations, with deviation from the expected slope becoming increasingly pronounced at greater degrees of dilution. CVs for the corrected sample concentrations across the six highest concentration dilutions were 5.9% and 21.3% for non-extracted and extracted samples, respectively. Parallelism with the Cayman Chemical kit was good across the full range of dilutions for both non-extracted (CV = 12.4%) and extracted saliva samples (CV = 11.7%; Figure 2). For AVP, both non-extracted (15.4%) and extracted (CV = 8.7%) saliva samples yielded good parallelism, although the range of values was greatly restricted, and near the lowest part of the standard curve for extracted samples.

**Figure 1.**
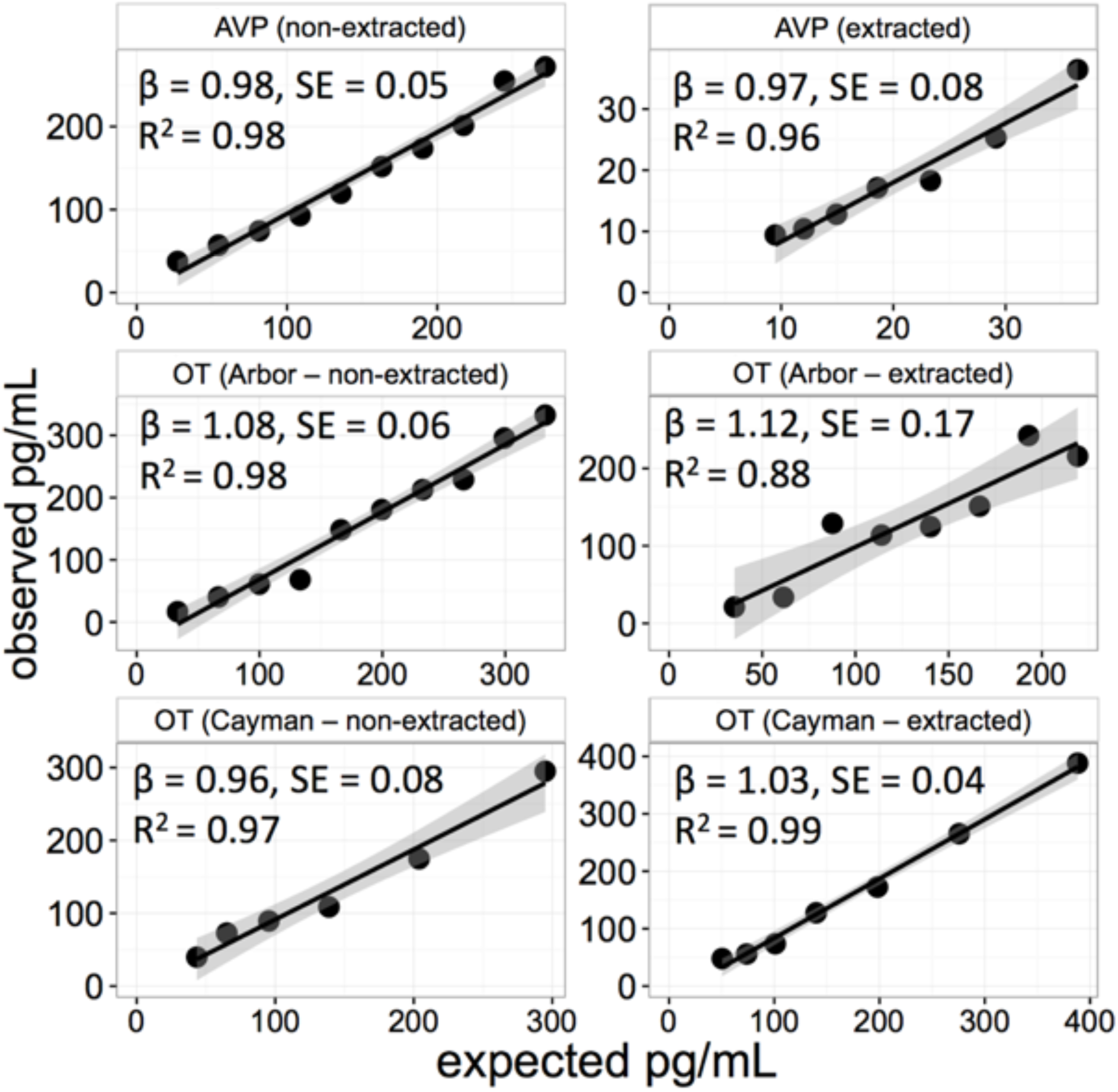
Regression fits predicting observed oxytocin (OT) and vasopressin (AVP) concentrations as a function of the expected concentration across a series of dilutions. β values indicate the slope, and SE, the standard error, of the regression fit which has an expected slope of 1. Shaded regions indicate the 95% confidence intervals for the slopes. Arbor: ELISA kit from Arbor Assays. Cayman: ELISA kit from Cayman Chemical.

**Figure 2.**
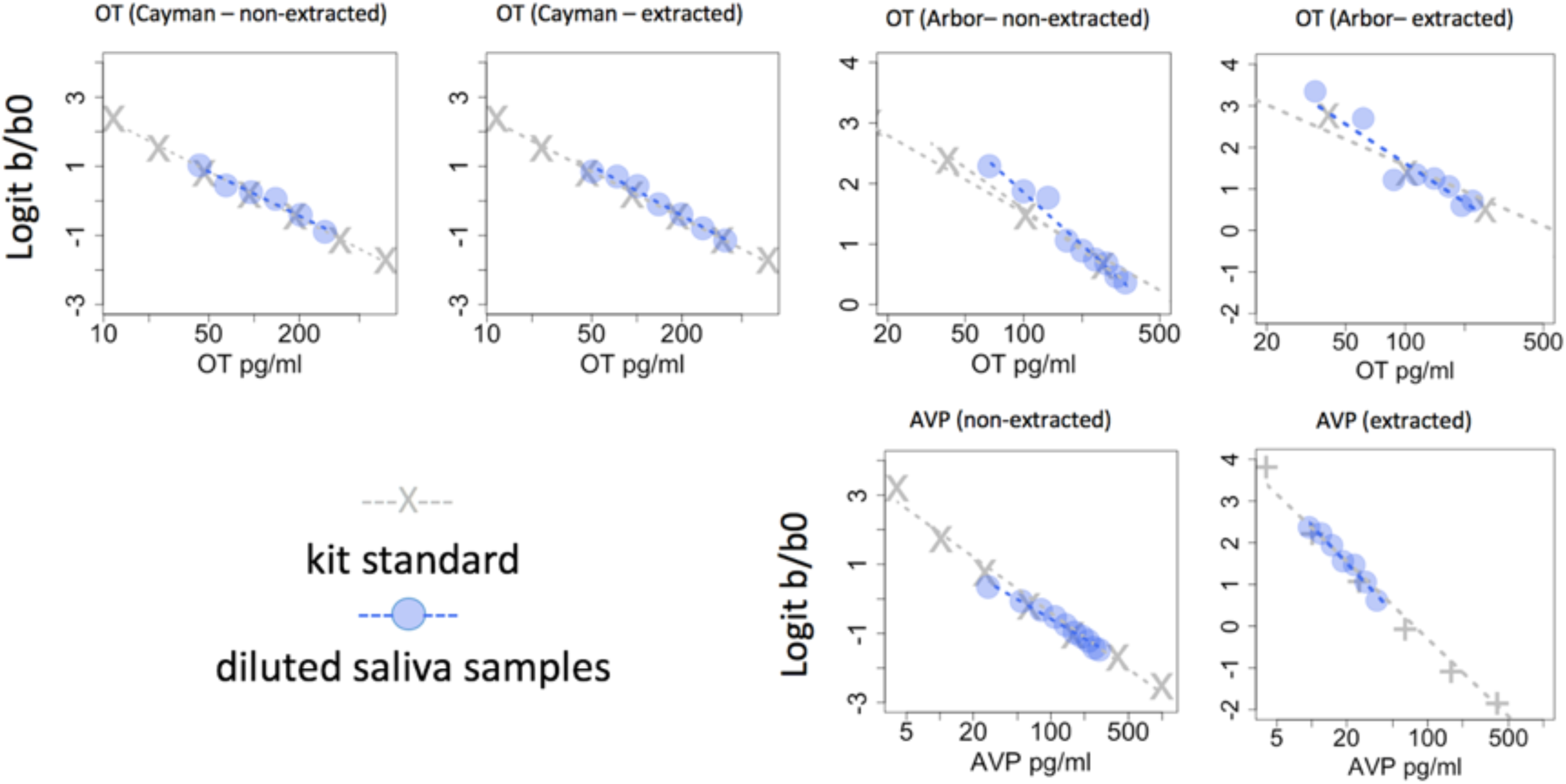
Parallelism for dilutions of non-extracted and extracted dog saliva samples. Gray X’s indicate kit standards and the gray dashed line reflects the linear relationship between binding and standard concentrations. Blue circles indicate different dilutions of pooled saliva samples. The blue dashed line shows a linear model predicting the expected sample concentration as a function of its binding at each dilution. Some samples near the lower limits of detection had clearly outlying values and were removed prior to analysis (see main text). Arbor: ELISA kit from Arbor Assays. Cayman: ELISA kit from Cayman Chemical.

Spike and recovery results with extracted and non-extracted saliva samples are shown in Table 1. For AVP, spike recoveries using an extracted pool of saliva were somewhat lower than expected, whereas recoveries with non-extracted saliva were closer to the expected value. Using the Arbor Assays kit for measurement of OT, we observed somewhat higher than expected recoveries using both extracted and non-extracted saliva samples. With the Cayman Chemical kit, recoveries were close to the expected values using both extracted and non-extracted saliva. Intra-assay CVs for non-extracted samples were 5.8% for OT (N = 10, mean = 825 pg/mL; Cayman Chemical) and 7.6% for AVP (N = 10, mean = 255 pg/mL). Inter-assay CVs were 15.9% for OT (N = 8, mean = 764 pg/mL; Cayman Chemical) and 4.6% for AVP (N = 7, mean = 520 pg/mL).

**Table 1.**
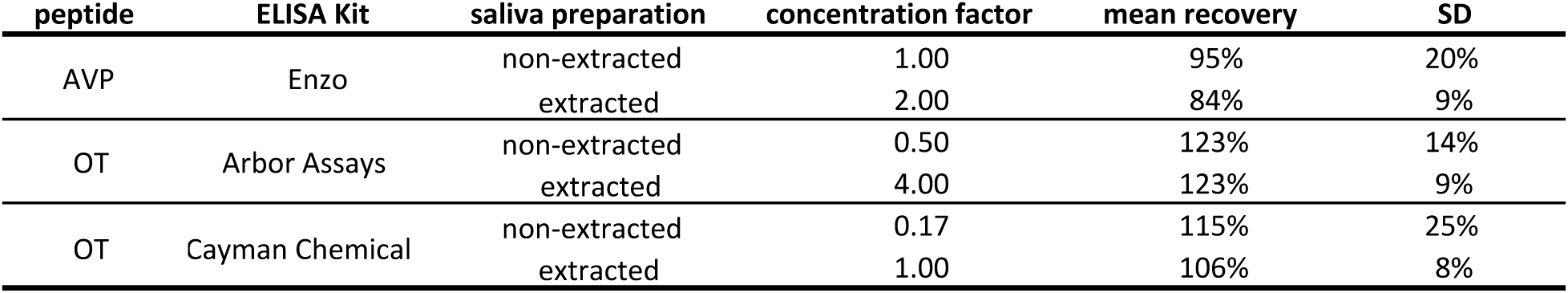
Results from recovery after spiking experiment. OT: Oxytocin, AVP: Arginine Vasopressin, SD = standard deviation of recovery.

| peptide | ELISA Kit | saliva preparation | concentration factor | mean recovery | SD |
| --- | --- | --- | --- | --- | --- |
| AVP | Enzo | non-extracted | 1.00 | 95% | 20% |
|  |  | extracted | 2.00 | 84% | 9% |
| OT | Arbor Assays | non-extracted | 0.50 | 123% | 14% |
|  |  | extracted | 4.00 | 123% | 9% |
| OT | Cayman Chemical | non-extracted | 0.17 | 115% | 25% |
|  |  | extracted | 1.00 | 106% | 8% |

### Sample Collection Techniques

Comparison of samples collected from the same dogs (each collected on a different day) using different collection procedures revealed variance accounted for by swab type (Children’s Swab or Salivette®), feeding immediately prior to the sample, and the use of citric acid to stimulate salivary flow (Figure 3; OT: F_3,53.4_ = 56.77, p < 0.01, AVP: F_3,56.5_ = 33.7, p < 0.01). For OT, Tukey HSD tests revealed that the Salivette® yielded OT concentrations significantly lower than the Children’s Swab (z = 8.00, p < 0.01), and these values were not correlated (r = -0.24, p = 0.33). Both citric acid and eating immediately prior to the sample (both samples collected using the Children’s Swab) tended to increase OT values, but only the food condition differed significantly from the baseline measure (citric acid vs. baseline: z = 1.77, p = 0.28; food vs. baseline: z = 5.28, p < 0.01). OT concentrations in samples collected following citric acid were strongly correlated with baseline measures collected on a different day from the same dogs (r = 0.57, p = 0.01). For samples collected following food, the correlation with baseline was positive, but not significant (r = 0.36, p = 0.21)

**Figure 3.**
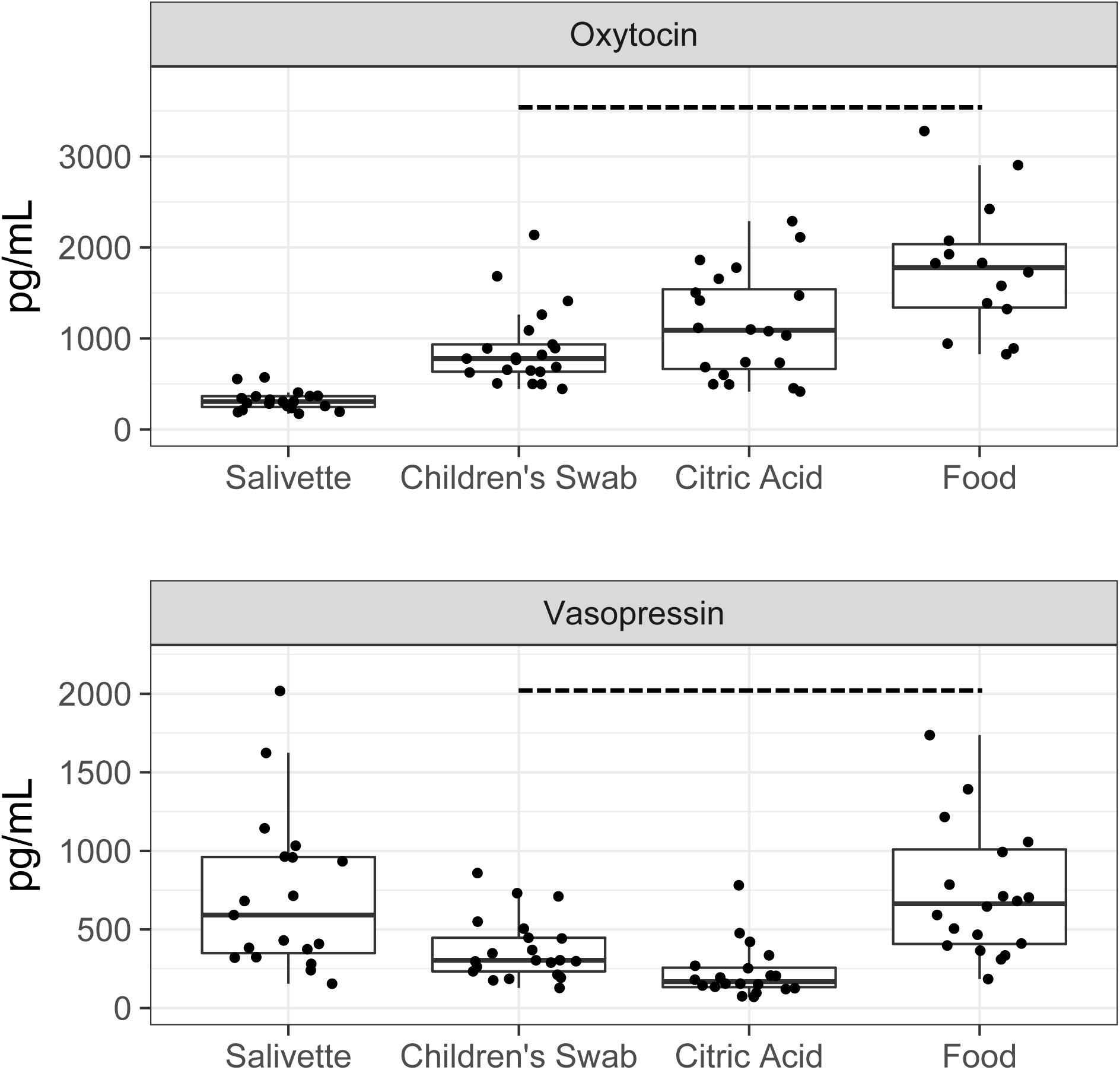
Oxytocin and vasopressin concentrations in saliva samples from the same dogs collected using different swabs (SalivaBio Children’s Swab, Sarstedt Salivette® – Cortisol), immediately following food consumption, and after salivary stimulation with citric acid. The dashed horizontal lines indicate sample groups collected with the Children’s Swab. The upper and lower bounds of the boxes correspond to the 25^th^ and 75^th^ percentiles of the data, and the whiskers above and below the boxes show the range of points within 1.5 times the interquartile range.

In contrast to OT, samples collected with the Salivette® yielded AVP concentrations that were significantly higher than samples collected with the Children’s Swab (Figure 3; z = -3.73, p < 0.01), but these values were positively correlated (r = 0.45, p = 0.05). Samples collected following citric acid yielded AVP concentrations that were significantly lower than baseline measures (z = -4.28, p < 0.01), and there was limited correlation between these measures (r = 0.31, p = 0.18). Lastly, as with OT, samples collected immediately following eating had higher AVP concentrations (z = 4.99, p < 0.01), but these values were nonetheless positively correlated with baseline measures (R = 0.59, p = 0.01).

### Biological Validation

Plasma oxytocin increased an average of 46.4% (SEM = 24%) and salivary oxytocin increased an average of 69.3% (SEM = 17.4%) from baseline to nursing (Figure 4). Although we detected a large increase in both matrices associated with nursing, the increase was statistically significant only for saliva (plasma: χ^2^ = 3.25, df = 1, p = 0.07; saliva: χ^2^ = 10.72, df = 1, p < 0.01). For the two Dams from whom we had post-nursing samples, both plasma and salivary OT decreased to approximately baseline levels shortly following nursing.

**Figure 4.**
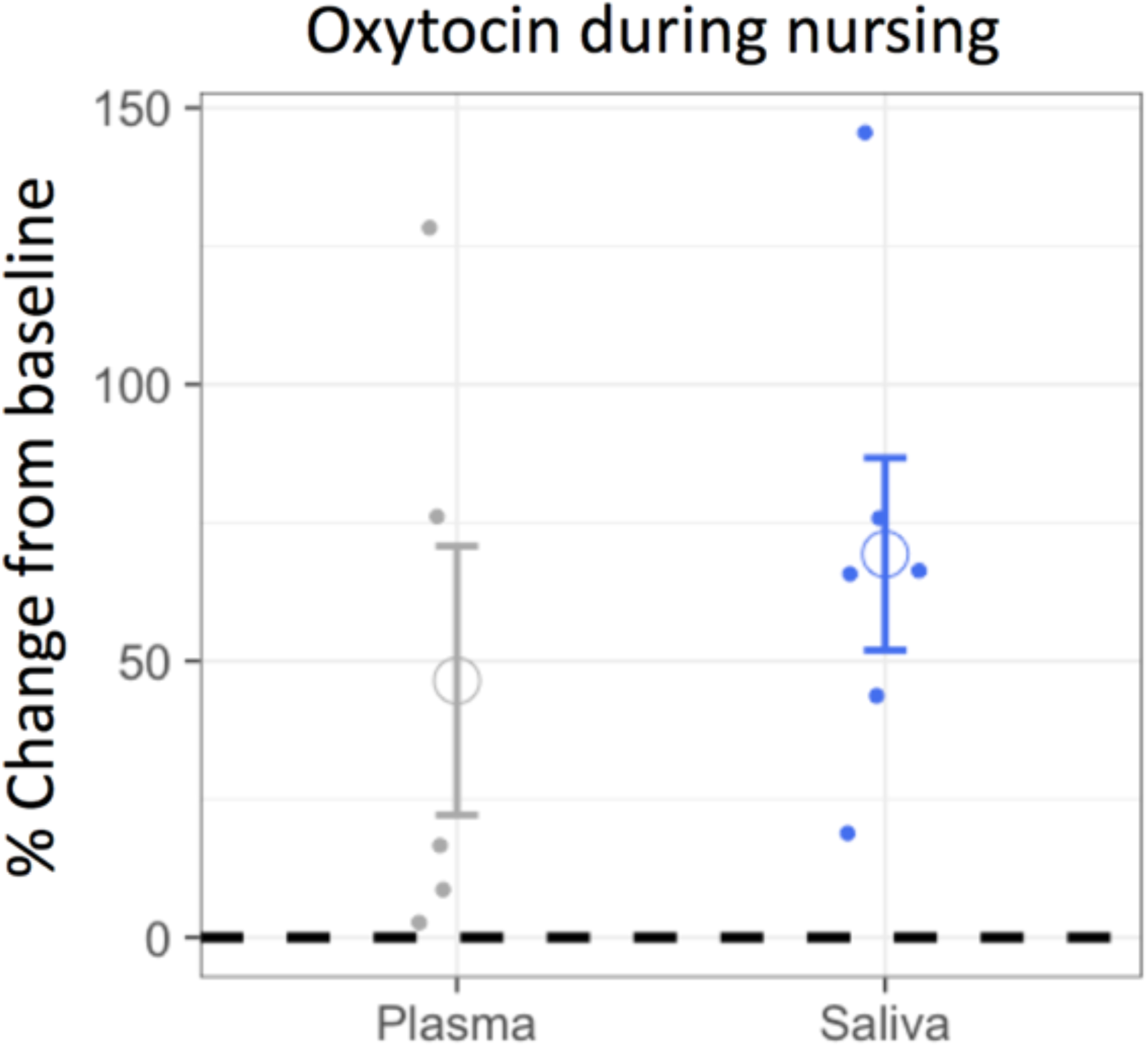
The percent change, from baseline, in plasma and salivary oxytocin during nursing. Solid points reflect the raw data and the open circles represent the mean change in each medium. Error bars reflect the standard error of the mean and the dashed horizontal line shows the null expectation of no change from baseline.

## Discussion

Our findings reveal that OT and AVP are present in dog saliva, that ELISAs using dog salivary samples exhibit good linearity, parallelism and spike recovery, and that salivary OT increases in parallel with plasma OT during lactation. Collectively these results suggest that salivary OT and AVP are promising biomarkers in domestic dogs, which have potential benefits related to both welfare, and research methodology. However, several aspects of our findings are surprising relative to previous studies of salivary OT/AVP in humans, and raise important questions about the biological processes through which these peptides reach saliva, as well as the range of techniques used for their detection.

We detected OT and AVP in dog saliva using multiple different ELISA kits, as well as HPLC-MS. However, the concentrations of OT and AVP detected by ELISA were much higher than those for HPLC-MS; moreover, for OT, the different ELISA kits yielded values in different ranges. These differences may be accounted for by a variety of factors. First, HPLC-MS detects a specific compound based on a known mass-to-charge ratio. In contrast, ELISAs may recognize not only the primary form of the target analyte, but also structurally related molecules, including precursor forms and biologically-related metabolites. Similarly, due to detection based on an antibody rather than a mass-to-charge ratio, ELISA may also detect the target analyte when it is bound to other components of the matrix. For these reasons, concentrations detected by immunoassay often exceed those from HPLC-MS (McCann, Gillingwater, & Keevil, 2005; Wood et al., 2008). Given that OT and AVP are characterized by multiple forms (Altstein & Gainer, 1988; Gainer, Altstein, & Whitnall, 1987; Green et al., 2001) and commonly bind to other molecules in biological matrices (Brandtzaeg et al., 2016; Martin & Carter, 2013), these phenomena may partially explain the differences we observed between the two ELISA kits, as well as between ELISA and HPLC-MS. Although the increased specificity of HPLC-MS comes with obvious advantages for interpretation of what is being measured, less specific detection methods may capture important biologically-related phenomena more broadly. For example, Galeandro et al. (2014) measured urinary corticoids in dogs with and without hypercortisolism using five different immunoassays in addition to gas-chromatography-mass spectrometry (GC-MS). Although GC-MS yielded significantly lower values than the immunoassays, it was no better at predicting disease status. Moreover, within the immunoassays, the least specific antibodies yielded the highest diagnostic accuracy (Galeandro et al., 2014).

Notably, the OT concentrations detected in dog saliva were dramatically higher than those in studies with humans. For example, in the validation study by White-Traut et al. (2009) salivary samples required a four-fold concentration to yield results in a reliable region of the standard curve, and corrected salivary OT values ranged from 6-61 pg/mL. Similar results have been reported in several other laboratories, and mean OT concentrations in human saliva are frequently reported to be < 5 pg/mL (Blagrove et al., 2012; de Jong et al., 2015; Feldman, Gordon, & Zagoory-Sharon, 2011; Grewen, Davenport, & Light, 2010; Holt-Lunstad, Birmingham, & Light, 2011; Javor et al., 2014). In contrast, dog salivary OT averaged 281 pg/mL and 694 pg/mL with the Arbor Assays and Cayman Chemical kits, respectively. Importantly, analysis of the same samples with and without extraction yielded highly correlated values in both kits (although extracted samples yielded considerably lower concentrations). However, even following extraction we obtained values much higher than those reported for non-extracted human saliva. Although we have not performed thorough validation studies with the OT ELISA kit from Enzo Life Sciences (the most commonly used kit for measurement of human salivary OT), pilot assays using that kit with non-extracted saliva from 16 dogs yielded similarly high values (mean = 690 pg/mL). Therefore, the drastically different results from humans and dogs are unlikely to be accounted for by variance between kit antibodies, or the presence of interfering substances that necessitate purification through solid-phase extraction. Given that recent proteomic studies have revealed hundreds of protein families in dog saliva which are not present in humans, as well as proteins in human saliva which are not present in dogs (Sousa-Pereira et al., 2015), it is possible that these factors relate to the observed species differences in OT concentrations.

In contrast to salivary OT, only non-extracted samples had salivary AVP concentrations in an acceptable range of the standard curve. In addition, AVP concentrations from the same samples analyzed with and without extraction were not correlated. Notably, the AVP concentrations detected by ELISA in extracted samples were frequently well below the value determined by HPLC-MS, suggesting that the extraction procedure may have eliminated substantial amounts of AVP prior to measurement. In contrast, AVP concentrations measured by ELISA in non-extracted samples were much higher than values determined by HPLC-MS, again suggesting the presence of structurally-related molecules in addition the primary form of free AVP. Nonetheless, across a range of dilutions, non-extracted salivary AVP samples yielded excellent linearity and parallelism, suggesting similar binding patterns between the AVP-immunoreactive compounds in dog saliva, and AVP standards.

To evaluate the effect of different saliva collection devices, we compared samples collected from the same dogs using the SalivaBio Children’s Swab and the Sarstedt Salivette® – Cortisol. Measures of both OT and AVP differed as a function of swab type but the direction of the effect was different between peptides. Specifically, OT concentrations were systematically higher in samples collected with the Children’s Swab, but AVP concentrations were significantly lower. Although both swabs are constructed of a synthetic material, whereas the Children’s Swab is designed to passively absorb saliva, the Salivette® has a larger diameter and is designed to be chewed on by a human participant. Therefore, the different physical properties of these swabs may affect which components of saliva are most readily absorbed, ultimately affecting concentrations in the sample. However, given the high structural similarity between OT and AVP, we do not know why different absorption profiles for the two swabs would lead to divergent effects between peptides. At present the Children’s Swab has been validated for use with at least 12 different analytes (e.g. cortisol, c-reactive protein, melatonin, alpha-amylase), whereas the Salivette® was developed and validated exclusively for cortisol measurement. Due its ease of use with dogs, and suitability for many salivary analytes, the Children’s Swab provides several advantages, however the differences between various swab types warrants further investigation.

The use of citric acid to stimulate salivary flow, or the consumption of food immediately before sample collection also affected OT and AVP measurements. Whereas citric acid led to modestly higher OT values, it had an opposite effect on AVP values. These findings are unlikely to be accounted for by changes to the salivary flow rate, in which case we would have expected decreased peptide concentrations in both cases. Instead, it is more likely that citric acid influences OT/AVP degradation in the sample (e.g. through a change in pH), or interferes with assay-specific reagents. Unlike citric acid, the effect of feeding before sample collection was similar for both peptides, leading to increased OT and AVP concentrations. Although feeding has been shown to increase OT in previous studies (Mitsui et al., 2011), and similarly may have effects on AVP, this is unlikely to explain these findings due to the very short time period (~30s) between feeding and sample collection. Despite the effects of feeding or salivary stimulation, OT and AVP concentrations in these samples tended to be correlated with baseline samples from the same dogs on a different day, suggesting individual stability in salivary peptide concentrations across time. However, given that both food and citric acid have the potential to affect OT/AVP concentrations, we recommend that researchers avoid these potential sources of interference.

In addition to providing a methodological validation, our data also suggest that measures of salivary OT respond to biological processes known to involve OT release. Specifically, we observed concurrent increases in salivary and plasma OT associated with the onset of milk letdown, followed by a decline to baseline levels following nursing (in two individuals for whom a third time point was measured). The relatively rapid changes in salivary OT contrast with findings for other hormones that reach peak concentrations in saliva ~10 minutes after those in blood (Hernandez et al., 2014). However, recent studies have revealed robust and similarly rapid rises in human salivary OT following the Regensburg Oxytocin Challenge (de Jong et al., 2015), which precede changes in salivary cortisol (presumably triggered by the same stimuli). Similarly, previous studies of salivary OT in nursing women revealed the highest OT concentrations immediately prior to nursing (White-Traut et al., 2009). This effect has been interpreted as reflecting an anticipatory rise in salivary OT in humans (White-Traut et al., 2009), as has been shown in other studies measuring plasma OT in nursing mothers (McNeilly, Robinson, Houston, & Howie, 1983). However, in our procedure dams were physically separated from their puppies prior to nursing, with no cues that nursing was imminent. Nonetheless, following the reunion of dams with their puppies, it typically took several minutes for dams to settle into a nursing position and for all puppies to secure a teat. Therefore, it is possible that the observed elevations in salivary OT partially reflect a rapid anticipatory response. However, the short time course now observed in several studies – coupled with the fact that OT and AVP have molecular weights approximately twice those of other hormones commonly measured in saliva (Horvat-Gordon et al., 2005) – present challenging, and as-of-yet unanswered questions about the mechanisms through which these hormones reach saliva.

At a practical level our findings have the potential to lead to improvements in welfare and research methodology in future studies of OT and AVP in dogs. Previously, studies measuring short-term changes in dog OT/AVP have relied on blood draws, which have greater potential to induce stress or momentary pain, and which are typically collected only in laboratory settings. In contrast, salivary samples are noninvasive and can be collected in diverse settings ranging from the laboratory to a dog park or private home. Compared to urine sampling, which is also noninvasive – salivary samples have the advantage of being collectable at short repeated intervals, without requiring an active response (i.e. urination) from the animal. Thus, the combination of saliva and urine sampling provides a flexible and noninvasive toolkit for assessing both short- and longer-term peptidergic activity in dogs. Lastly, our findings reveal that dog salivary OT samples do not require an extraction procedure, eliminating a costly and time-intensive component of sample processing. Therefore, the measures described herein provide researchers with a new set of noninvasive and methodologically-sound tools for investigating the roles of OT and AVP in dog behavior and cognition, as well as human-animal interaction.

## Acknowledgements

We are grateful to Brenda Kennedy and staff at Canine Companions for Independence for their help with this research. We thank Hossein Nazarloo, Martina Heer, and Taichi Inui for helpful conversations, and Susan Alberts for allowing us to work her laboratory. We gratefully acknowledge support from the WALTHAM® Centre for Pet Nutrition, which funded this research.

